# Facial expressions in mice reveal latent cognitive variables and their neural correlates

**DOI:** 10.1101/2024.06.02.597005

**Authors:** Fanny Cazettes, Davide Reato, Elisabete Augusto, Alfonso Renart, Zachary F. Mainen

**Affiliations:** Champalimaud Foundation; Av. Brasília, 1400-038 Lisboa, Portugal; Institut de Neurosciences de la Timone, CNRS and Aix Marseille Université; 27 Bd Jean Moulin, 13005 Marseille; Mines Saint-Etienne; Centre CMP, Departement BEL, F - 13541 Gardanne, France

**Author notes:** ^†^Equal contribution. ^‡^Equal contribution.

## Abstract

Brain activity controls adaptive behavior but also drives unintentional incidental movements. Such movements could thus potentially be used to read out internal cognitive variables also neurally computed. Establishing this, however, would require ruling out that incidental movements reflect cognition only because they are coupled with task-related responses through the biomechanics of the body. We addressed this issue in a foraging task for mice where multiple decision variables are simultaneously encoded even if, at any given time, only one of them is used. We found that characteristic features of the face simultaneously encode not only the currently used decision variables, but also independent and unexpressed ones, and we show that these features partially originate from neural activity in the secondary motor cortex. Our results suggest that the face reflects ongoing computations above and beyond those related to task demands, demonstrating the ability of noninvasive monitoring to expose otherwise latent cognitive states.

## INTRODUCTION

The brain is tasked with controlling the body through extensive nerve pathways that activate muscles. Patterns of motor activity are informed by brain computations which depend on both the current states of the environment and the organism, as well as the organism’s past experience. In attempting to unravel the mechanisms of behavior, it is common to assume that movements are adaptive output meant to act upon the world to reach desired goals. However, this premise has an important caveat. It is long known that the body expresses a variety of movements that are neither intentional nor obviously adaptive. Rather, they can be seen as an unintentional and/or unconscious “leakage” of brain activity into the body. The importance of “incidental” facial expressions and “body language” has long been highlighted in popular psychology. For instance, video of the face can be used to detect a large array of nuanced emotional expressions (*1*), and incidental features of mobile phone interaction can be used to diagnose psychiatric conditions (*2*). While these kinds of studies clearly attest to the potential information latent in bodily expressions, concomitant measurements of neural variables are requisite to better understand how bodily expressions reveal internal cognitive processes.

Leakage of cognitive operations into the body was documented in studies showing that posture reflects the contents of working memory in rats (*3*, *4*) as well as in experiments showing that reflex gains in the arm reflect accumulated evidence during perceptual decision-making in humans (*5*). In these examples, the influence of cognitive variables on observable behavior is related to the action being used to execute the task at hand. However, not all internal states are directly related to task execution, and may serve various other purposes (*6–8*). Distinct multidimensional facial expressions can be evoked by salient, behaviorally-relevant stimuli in mice (*9*). While emotional expressions have traditionally been restricted to a small canonical set (e.g., pleasure, disgust, fear, surprise, etc.), empirical approaches suggest a much higher dimensionality (*1*). Similarly, pupil diameter is linked not only to changes in luminance, directly relevant for seeing, but also to more abstract cognitive variables related to arousal and uncertainty, even when they have no direct relationship to perception (*10–12*). In line with this, in mice, even neutral sounds of different spectral or temporal content evoke stimulus-specific facial movements (*13*, *14*).

To make the best case for the link between internal state and bodily expressions, rich behavioral observations using high-resolution video recordings during task execution can be combined with large-scale neural recordings, as pioneered by recent work in mice and non-human primates (*15–20*). These studies have shown that incidental movements can account for a larger fraction of the variance of neural activity than task-related movements (but see (*20*)), and are informative about the internal state of engagement (see also (*21*)), and that the extent to which incidental movements shape neural activity depends on the brain area (*18*, *20*).

Experiments of this sort, conducted in a task together with neural recordings, can reveal precise relationships between computational variables and both neural and bodily expressions. However, they face a critical challenge: it is difficult to rule out that incidental movements become meaningful, i.e., associated with specific cognitive variables, simply because physical constraints of the body link them to other movements that are necessary to report these variables in the first place (*18*). To address whether incidental movements are only linked to ongoing computations in this ‘trivial’ way, it would be necessary to have a way of defining meaning independently of the current behavioral strategy used to solve the task. Recently, (*22*) showed that during the performance of a foraging task, an entire family of different foraging-related decision variables can be decoded from the brain in a sustained fashion throughout a behavioral session. This family of decision variables is a collection of complex and dynamic internal state markers computed by applying different algorithms to information being accumulated over relatively long time scales (several seconds). During task performance, one of these decision variables primarily manifests itself in the mouse’s foraging choices. The rest of the variables in the family, which are equally complex, are computed neurally and are decodable throughout, but are *not* expressed in the task performance of the mouse at that particular time. Instead, they may be expressed at other times during the session. This provides an ideal situation to ask whether meaningful but currently task-unrelated internal states (decision variables) can be expressed incidentally in the face or body. If existing, this relationship between cognitive variables and movement would, by definition, avoid being trivially explainable through the demands of the task report.

Here, we monitored mice’s facial movements while simultaneously recording or manipulating the activity of large ensembles of neurons in frontal cortical regions during the foraging task of Cazettes’s et al. This allowed us to directly compare the ability of facial expressions to that of large-scale neural recordings to reveal computationally well-defined but *task-irrelevant* internal state variables. We found that diverse decision variables with varying dynamics – even currently unused ones – could be linearly decoded from facial movements with a similar accuracy as neural recordings. Each decision variable correlated with characteristic facial features, consistently across mice, suggesting a stereotyped link between decision variables and facial expressions. Through correlational (recording) and causal (optogenetics) experiments, we demonstrated that the facial expressions reflecting the decision variables partially stemmed from neural activity in the secondary motor cortex (M2). These results illustrate approaches for decoding seemingly meaningless facial expressions to infer meaningful neural states, demonstrating the potential of non-invasive monitoring to reveal otherwise hidden cognitive activity.

## RESULTS

### Animals used multiple decision variables to time their foraging decisions

We analyzed data from a probabilistic foraging task in which mice could navigate between two virtual resource sites and collect drops of sugary water by freely licking at a port (Fig. 1A, but also see methods and (*22*) for details on the behavioral approach). By design, only one of the resource sites delivered rewards at any given time. A hidden state determined which of two sites provided rewards with a fixed probability (0.9) after a lick. The hidden state also had a fixed probability (0.3) of transitioning from rewarding to depleted after each lick, but never the other way around. Once a specific resource site was depleted, the only way to receive reward again was to reach the other resource site. The presence of the state transition thus required the animal to decide after every lick whether to stay at or to leave the current resource site.

**Figure 1.**
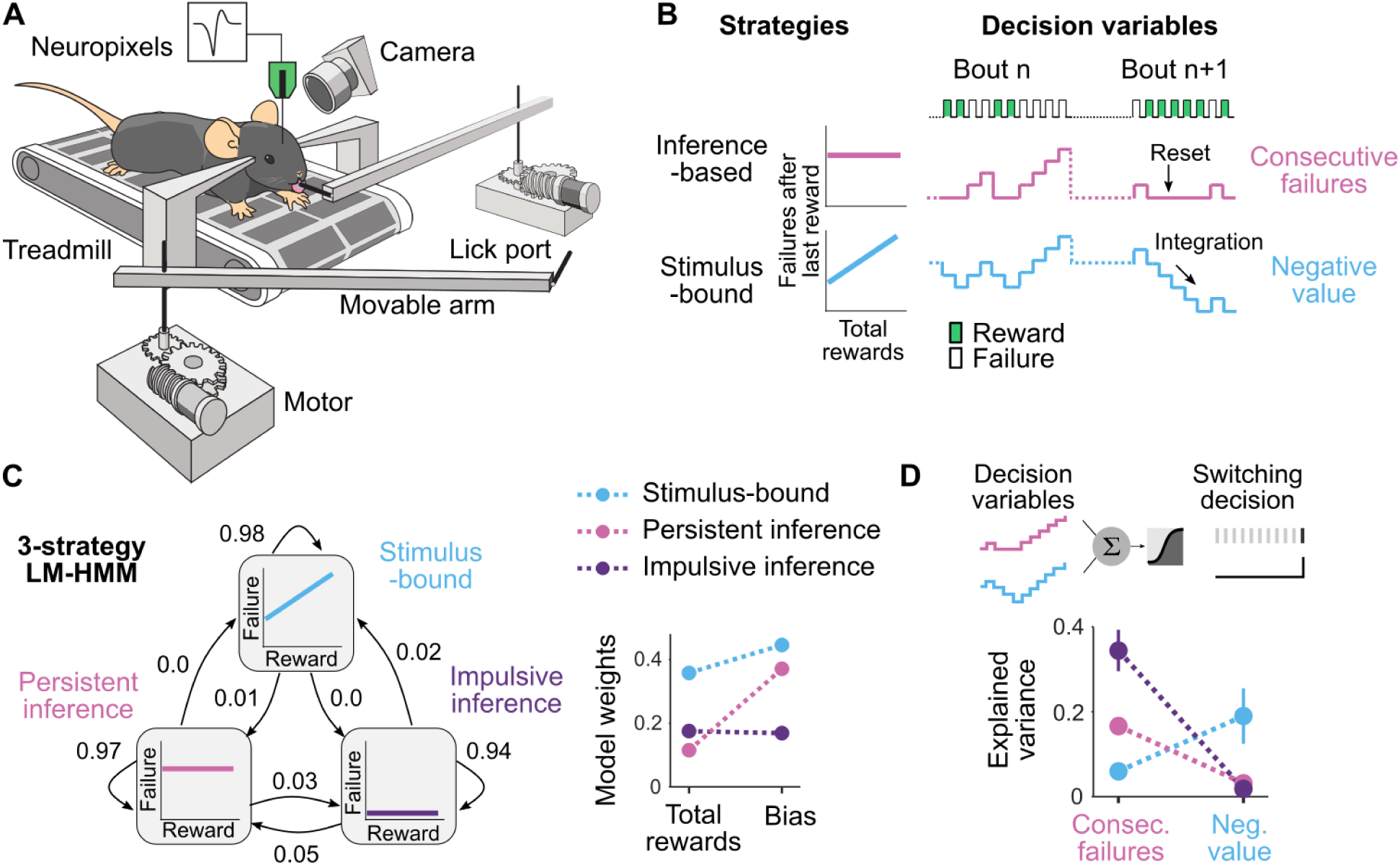
Task, strategies & decision variables. **A.** A head-fixed mouse is positioned on a treadmill and can forage at one of two sites (the two movable arms). Head-fixation enables the monitoring of facial movements using high-speed cameras (60 fps) and provides better stability for high-throughput electrophysiology (Neuropixels). The mouse can choose to switch between sites at any time by running a set distance on the treadmill. During site-switching, the front arm moves away, and the distal one moves into place. **B.** Different strategies for timing the decision to leave the site are based on distinct decision variables. The inference-based strategy (blue) is reward-independent and resets with each reward, while the stimulus-bound strategy (pink) depends on reward integration. **C.** Left: Illustration of the LM-HMM with three different strategies (labeled ‘Stimulus-bound,’ ‘Persistent inference,’ and ‘Impulsive inference’). High self-transition probabilities of 0.98, 0.97, and 0.94 indicate that strategies typically persist for many consecutive bouts. Transition probabilities are shown by the arrows between states. Right: LM weights for the three-strategy model fitted to all sessions simultaneously (N = 10). **D.** The variance explained by the two different decision variables during each strategy (mean ± s.d. across 10 iterations of the cross-validated logistic regression).

As described in detail in (*23*) and in (*22*), mice could decide to leave a foraging site based on multiple decision variables that relied on different combinations of observable events (i.e., rewarded and failed foraging attempts, Fig. 1B). In theory, the optimal solution to time the decision to leave is to infer the hidden state of the resource site by counting the number of consecutive failures, which are evidence that the site is depleted, with a complete reset upon reward delivery, which signals with certainty that rewards are still available at the site (Fig. 1B, pink). We refer to this reward-independent strategy as *inference-based*. Alternatively, another common strategy is to time the decision to leave based on a running estimate of how much reward is received at the site, which is equivalent to calculating the net negative value of the site (Fig. 1B, blue), staying longer the more rewards are delivered. We refer to this reward-dependent strategy as *stimulus-bound*.

In previous work, we showed that mice relied on these two strategies interchangeably across different sessions and even within the same sessions across different epochs (*22*). Using a framework based on hidden Markov models (HMM) combined with linear regression models (LM, Fig. 1C left), we fit the number of consecutive failures that the animal is willing to accept before switching sites. This was achieved using two inputs: (1) the total number of rewards, which allows a distinction between inference-based (i.e. reward independent) and stimulus-bound (i.e. reward dependent) strategies, as in Fig. 1B left, and (2) a constant bias, which reflects the level of impulsivity of the animal (notice that high bias means large number of failures accepted before switching sites, i.e., low impulsivity). Each hidden state in the model captures a specific dependence of consecutive failures on the total rewards and the bias, characterizing a particular decision-making strategy. Here, a model with three states best described the foraging decision and yielded interpretable and persistent strategies (Fig. 1C and (*22*)). One of the strategies had large weight on the number of rewards, indicative of a stimulus-bound strategy, while the other two had small weights on rewards, consistent with the inference (Fig. 1C right).

To estimate how the decision variables explained the mice behavior in each strategy, we used regularized logistic regressions to model the probability that each lick (n = 5877 licks in stimulus-bound, n = 6896 licks in persistent inference and n = 1593 licks in impulsive inference, across 10 sessions) was the last one in the bout, considering simultaneously the consecutive failure and the negative value decision variables as predictors (Fig. 1D top). This multivariate approach confirmed that the consecutive failures better explained the inference-based strategies, while the negative value better explained the stimulus bound strategy (Fig. 1D bottom).

### Decision variables are reflected in facial movements

To test whether fine movement of the face reflected latent decision variables, we extracted high-dimensional temporal representations of the movement signals by applying singular value decomposition to the motion energy movies (*24*) (Fig. 2A left). We extracted the principal components of the movements (movement PCs, N = 100) from different regions of interest: (1) the entire face, (2) the upper face, excluding the tongue area where most of the movement occurred during licking, and (3) the body parts visible in the field of view. The movement PCs were binned in 200 ms windows around each licking event (Fig. 2A middle), which directly correspond to discrete changes in the decision variables. We then predicted different decision variables on a trial-by-trial basis using cross-validated and regularized generalized linear models (GLMs, Fig. 2A right, see methods).

**Figure 2.**
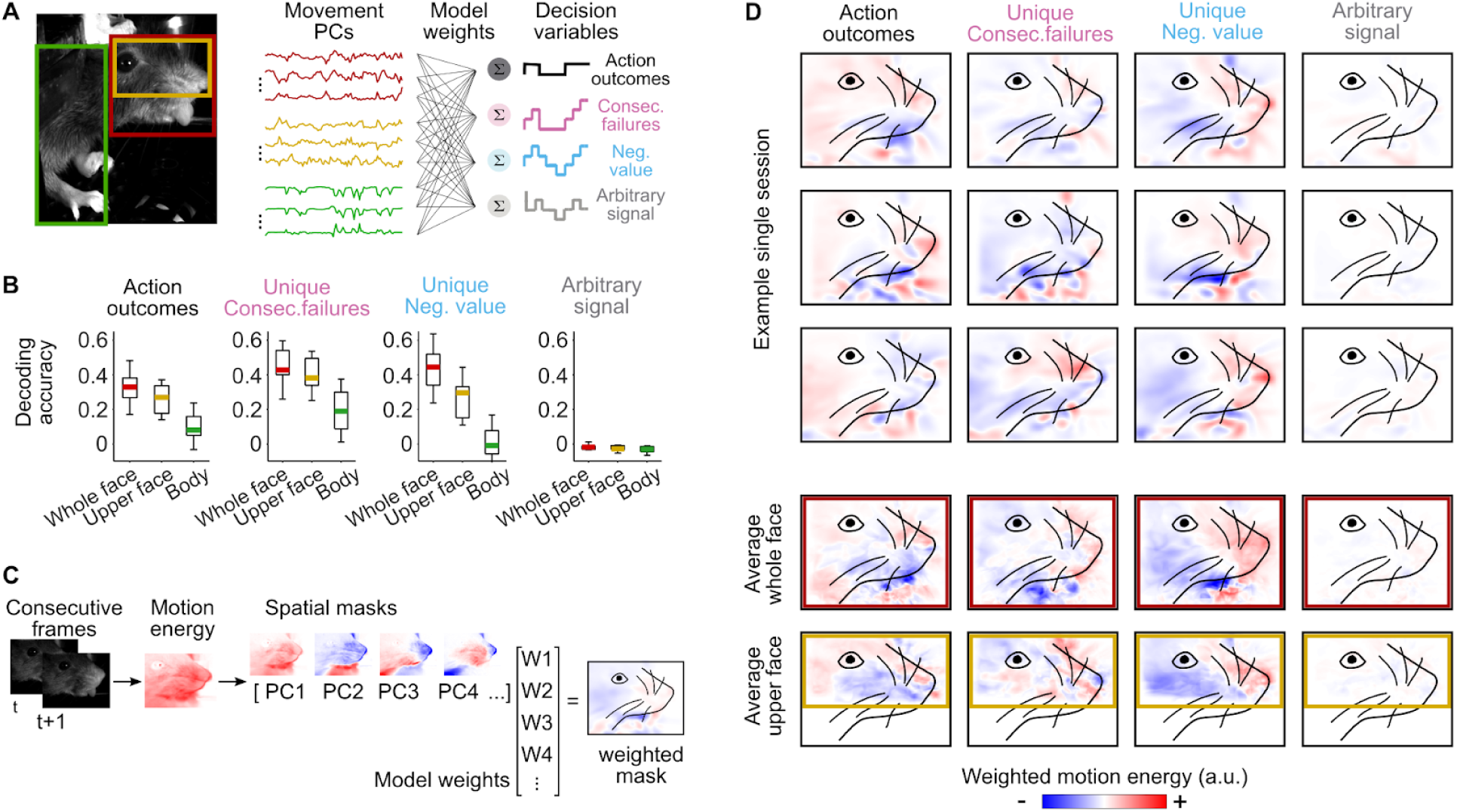
Stereotyped facial expressions of decision variables. **A.** Left: Example principal components extracted using SVD on the motion energy movie (Movement PCs) from three regions of interest. Right: The movement PCs are used as predictors in multivariate regression models to predict a set of decision variables and arbitrary signals. **B.** Decoding accuracy (cross-validated R2) in each region for different decision variables (median across 10 sessions and 25th and 75th percentiles; whiskers represent minimum and maximum values). **C.** After computing the motion energy as the absolute value of the difference of consecutive frames, facial masks corresponding to the facial movement PCs were weighted by the models’ weights. The weighted sum of the facial masks indicate the average motion energy for a given decision variable. In this example, we show the average motion energy for the negative value during a single session. **D.** Motion energy weighted by the models’ weights for single example sessions (top) and averaged across sessions (N = 10, middle for the whole face and bottom for the upper face). Red represents more movement than average, while blue indicates less movement than average.

To test whether facial movements reflect latent computations rather than solely responding to instantaneous stimuli, we orthogonalized the decision variables both to the observable action outcomes and among themselves, and attempted to decode the ‘unique’ aspect of each decision variable (i.e., the residual after regressing it to the others and to the action outcomes). As a control, we also tested the decoding of arbitrary signals possessing the same power spectrum as the decision variables. We found that while we could not decode the arbitrary signals from any movement PCs, we could decode action outcomes and decision variables with relatively high accuracy but mostly from facial movement PCs (Fig. 2B).

To test whether characteristic features of the face expressed different decision variables, we examined the averaged spatial distribution of facial motion weighted by the models’ weights (Fig. 2C). Noticeable differences in facial movement were associated with different decision variables consistently across mice (Fig. 2D, see also Fig. S3 for further evidence on an independent cohort of mice). For instance, the value corresponded to more movement around the nose than around the cheek while the consecutive failures corresponded to a more subtle pattern of facial movement. Altogether, these results suggest that decision variables are reflected in stereotypical facial expressions.

### Facial expression reflect a reservoir of decision variables

Because the mice transition between periods where they use one of the three identified strategies (Fig. 1C), and because behavior during each strategy is best predicted by different decision variables (Fig. 1D), the results in Fig. 2 are consistent with two scenarios. First, it could be that the characteristic patterns of facial motion become associated with each of the decision variables only because the mice move slightly differently during each of the behavioral strategies. As noted in the introduction, this corresponds to the ‘trivial’ case where cognitive variables leak into incidental movements only because there are different patterns of task-related movement associated with the report of those cognitive variables. Alternatively, the link between facial movement and decision variables could exist above and beyond the strategy currently used by the mouse. For instance, it could be that the face instantaneously reflects the stimulus-bound decision variable during periods where the mouse is solving the task using an inference-based decision variable. While this might seem surprising, in previous work we have shown that neuronal activity in the frontal cortex of mice retains information about multiple decision variables simultaneously, even when the variables do not explain the foraging decision of mice (*22*). Thus, we attempted to test if the face, like the neurons, contains information about particular decision variables even during periods where those decision variables are not guiding behavior.

To address this question, we registered and concatenated the videos to extract facial movement PCs across sessions (Fig. 3A, Fig. S1). This procedure allowed us to compare the same facial movement PCs in different strategies and their relative variance (Fig. 3B). From these facial movement PCs, we first asked whether we could also decode the different strategies. Using the same regularized and cross-validated multivariate regression approach as previously described, we could predict the probability of being in a given strategy from facial movement PCs (Fig. 3C). The critical test is then to decode the different decision variables during each strategy separately. If the decision variables are expressed on the face only when guiding behavior, the portion of the signal that is unique to each variable should only be decoded in the corresponding strategy (Fig. 3D, left). Alternatively, if the decision variables are expressed on the face even when not guiding behavior, the decoding accuracy should be independent of the strategy (Fig. 3D, right). We found that the decoding accuracy of the decision variables was unequivocally strategy-independent (Fig. 3E). These results suggest that, at least in the context of our task, when the brain calculates a given decision variable, that variable becomes reflected in the facial expressions of the mouse, even if that decision variable is not currently used to guide behavior (and is orthogonal to the alternative decision variables which do play this guiding role).

**Figure 3.**
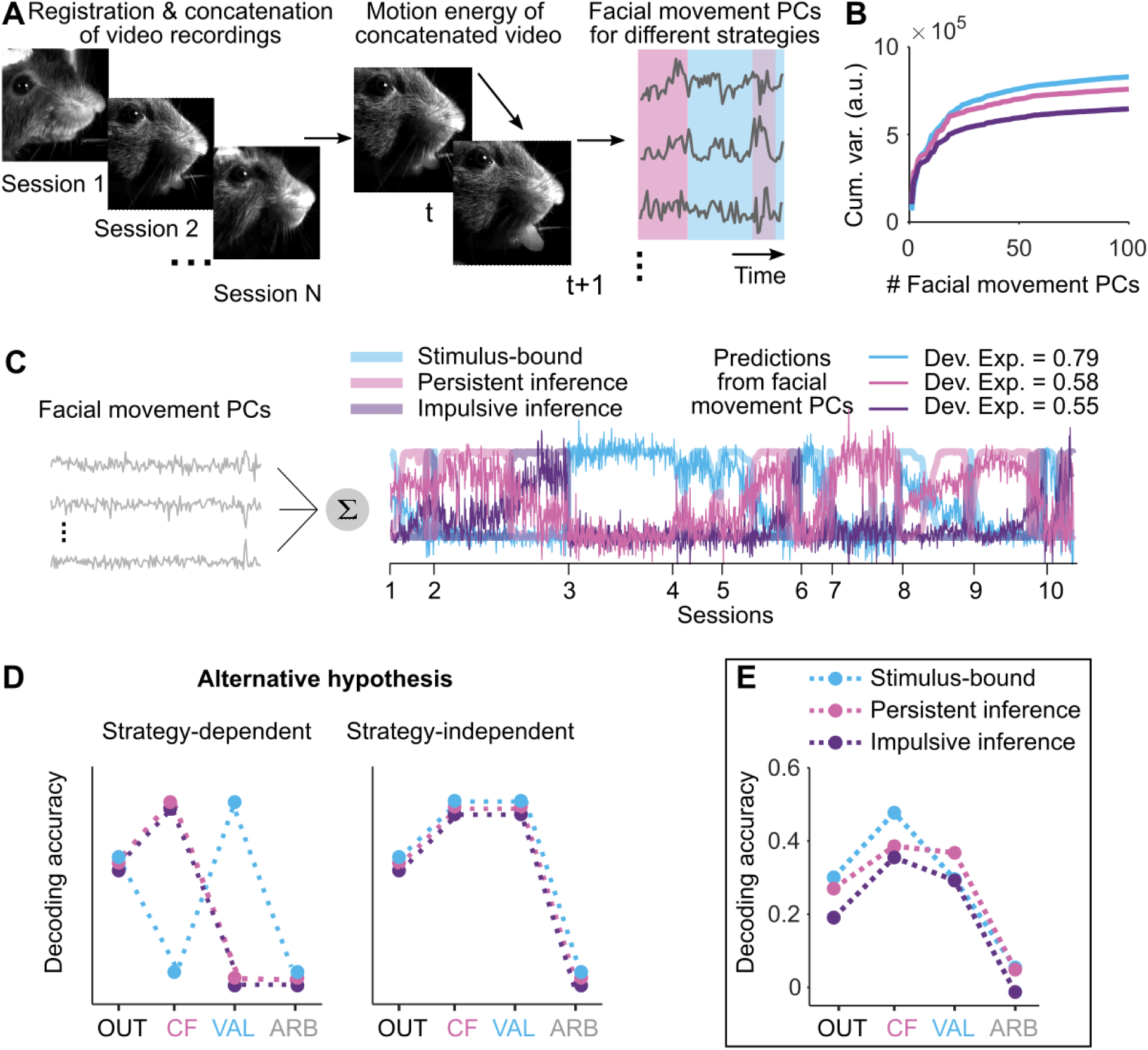
Facial expression of decision variables does not depend on the strategy. **A.** Registration and concatenation of video recordings to extract the movement PCs of the face across different strategies. **B.** Cumulative sum of the variance of the movement PCs in each strategy epoch. **C.** Illustration of the multivariate regression method for predicting the strategies (thick color traces) from the facial movement PCs (gray traces). The predictions of the models (thin color lines, which are the weighted sums of the facial movement PCs) track quite faithfully the probability of being in a given strategy (thick color traces). **D.** Schematic representation of hypothetical performance of strategy-dependent (left) vs. strategy independent (right) models. Left: the decoding accuracy of the decision variables depends on the strategy. Right: the decoding accuracy of the decision variables is independent of the strategy. **E.** The decoding accuracy of the decision variables from facial movement PCs is strategy independent (similar to right D).

### Comparisons of facial and neural expressions of decision variables

The capability to simultaneously decode multiple variables from facial motion, independently of strategy, mirrors the properties of the secondary motor cortex (M2), where we have previously shown the simultaneous decoding of multiple decision variables from neural activity (22). This raises the question of which of the two brings about the other. On the one hand, decision variables could manifest in neural activity as efference copies of motor signals. On the other hand, neurons could compute decision variables that are subsequently reflected in facial movements. If the latter is true, there should be ‘covert’ neural representations of decision variables in neural activity preceding the ‘overt’ expressions of decision variables in facial movement. To test this idea, we evaluated the accuracy and latency of representations of decision variables across different cortical regions (M2: N = 67 ± 26 neurons; the orbitofrontal cortex - OFC: N = 58 ± 18 neurons; the olfactory cortex - OC: N = 28 ± 13 neurons, median ± m.a.d. across 10 sessions; Fig. S2), comparing them to those observed in facial movements.

We generated, for each neuron and facial movement PC, time lag versions of the binned activity using a sliding window with increasing delays from the time of the lick (Fig. 4A). We then used each lagged series of neural or facial movement activity to independently predict the different decision variables with cross-validated and regularized GLMs. The representations of decision variables by each cortical region and facial movement PCs varied depending on the time delay, peaking at different accuracy levels (Fig. 4B). In particular, the accuracy from facial movement PCs was significantly greater than that of neural representations from all recorded regions (Fig. 4D,E; ΔAccuracy M2: 0.13 ± 0.1, p = 0.00004; ΔAccuracy OFC: 0.34 ± 0.1, p = 0.000002; Δlatency OC: 0.43 ± 0.08, p = 0.000002; median ± m.a.d. across 10 sessions, Wilcoxon signed rank test).

**Figure 4.**
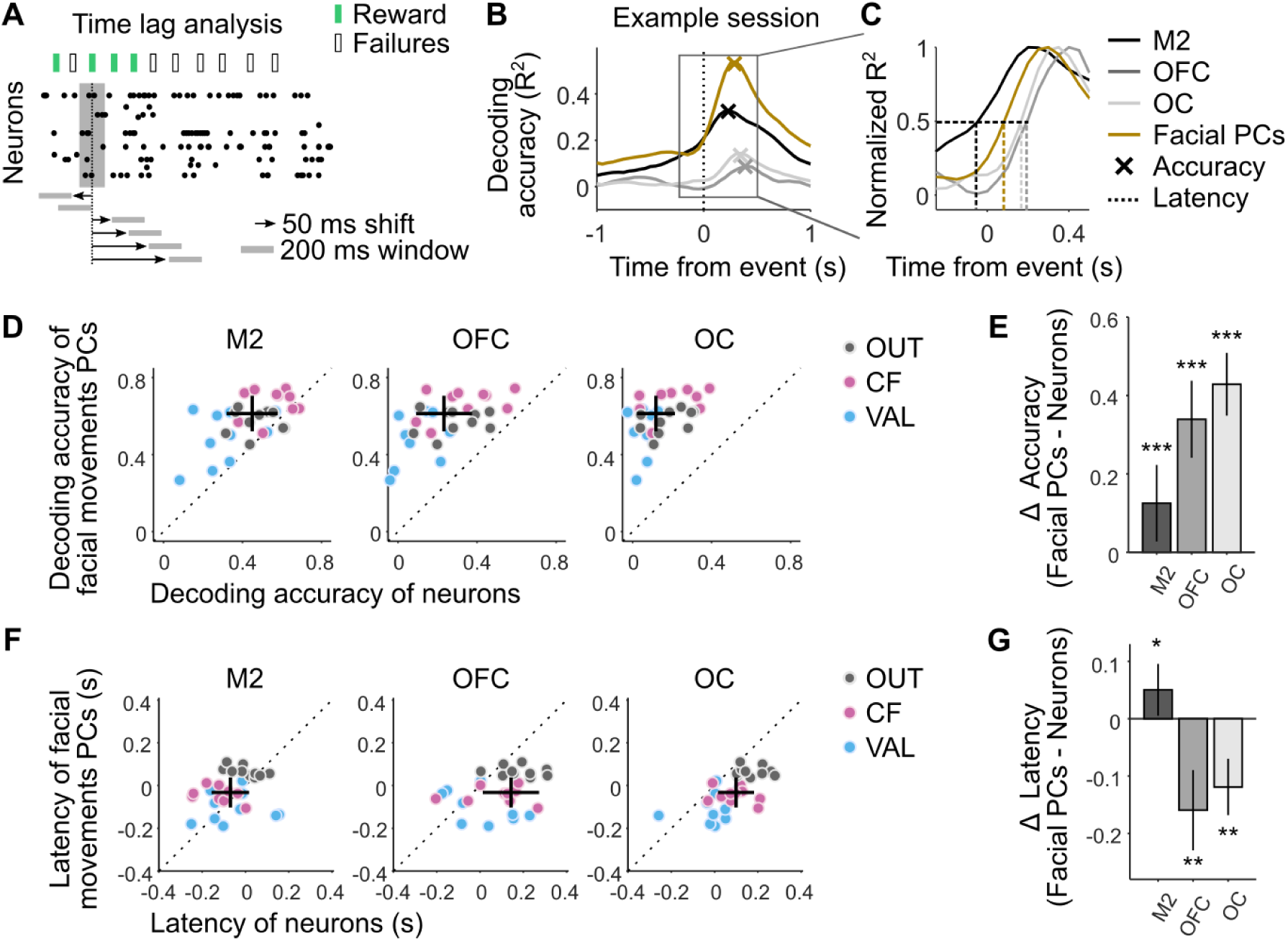
Expressions of decision variables: facial movement vs. neurons. **A.** A 200 ms sliding window with a 75% overlap was used to generate time-lag series of neural and movement activity. **B.** Example decoding accuracy (cross-validated R^2^) of multivariate regression predicting decision variables from facial movement PCs (gold) or neural activity in different brain regions (grays) as a function of the time delay of the predictors. The crosses indicate the maximum accuracy. **C.** Example estimation of latencies corresponding to the midpoint R^2^ relative to each brain region and facial movement PCs. **D.** Peak decoding accuracy from facial movement PCs against peak decoding accuracy from neurons for the different decision variables. Dots (1 gray, 1 pink, and 1 blue per session, N = 10 sessions) above the identity correspond to representations of decision variables from facial movement PCs being more accurate than that of neural representations. Black cross is the median ± m.a.d. across all data points. **E.** Difference between the accuracy from facial movement PCs and from neurons in each brain region (median ± m.a.d. across 10 sessions). Stars indicate significance of the Wilcoxon signed-rank test (p < 0.05: one star; p < 0.01: two stars; p < 0.001: three stars). **F.** Same as in (D) but for decoding latency. Dots above the identity correspond to representations of decision variables from facial movement PCs following neural representations. **G.** Same as in (E) but for decoding latency.

The latency of representations of decision variables, estimated from the time delay corresponding to the midpoint decoding accuracy, also varied across brain regions and facial movement PCs (Fig. 4C). We found that decision variables were decodable from the facial movements prior to OFC and OC (Fig. 4F, G; ΔLatency OFC: -0.16 ± 0.0059 s, p = 0.0059; ΔLatency OC: -0.12 ± 0.05 s, p = 0.002; median ± m.a.d., Wilcoxon signed rank test). Yet, representations of decision variables emerged after the neural representations in M2 (ΔLatency M2: 0.05 ± 0.05 s, p = 0.037; median ± m.a.d., Wilcoxon signed rank test). These results suggest that M2 may participate in the generation of the facial expressions of decision variables.

### Effect of M2 inactivation on facial expressions of decision variables

We have previously reported in (*22*) that the partial inactivation of M2 significantly decreased the predictive power of the decision variables for explaining the switching behavior, suggesting that M2 is part of the neural pathway through which the decision variables shape behavior during the foraging task. Here, the emergence of decision variable representations in M2 prior to their facial expression (Fig. 4F,G) suggests that M2 activity may also contribute to generating micro-movement related to decision variables. To determine whether M2 activity is required for the facial expression of decision variables, we performed bilateral optogenetic inactivation of M2 while video monitoring facial movements during the foraging task. Specifically, we used VGAT-ChR2 mice, which express the excitatory opsin channelrhodopsin-2 in inhibitory GABAergic neurons, to silence M2 in 30% of randomly selected behavioral bouts (Fig. 5A). We examined 48 sessions from 8 mice, 5 of which were ChR2-expressing and three of which were control wild-type littermates that express no inhibitory opsin implanted and stimulated in the same manner.

**Figure 5.**
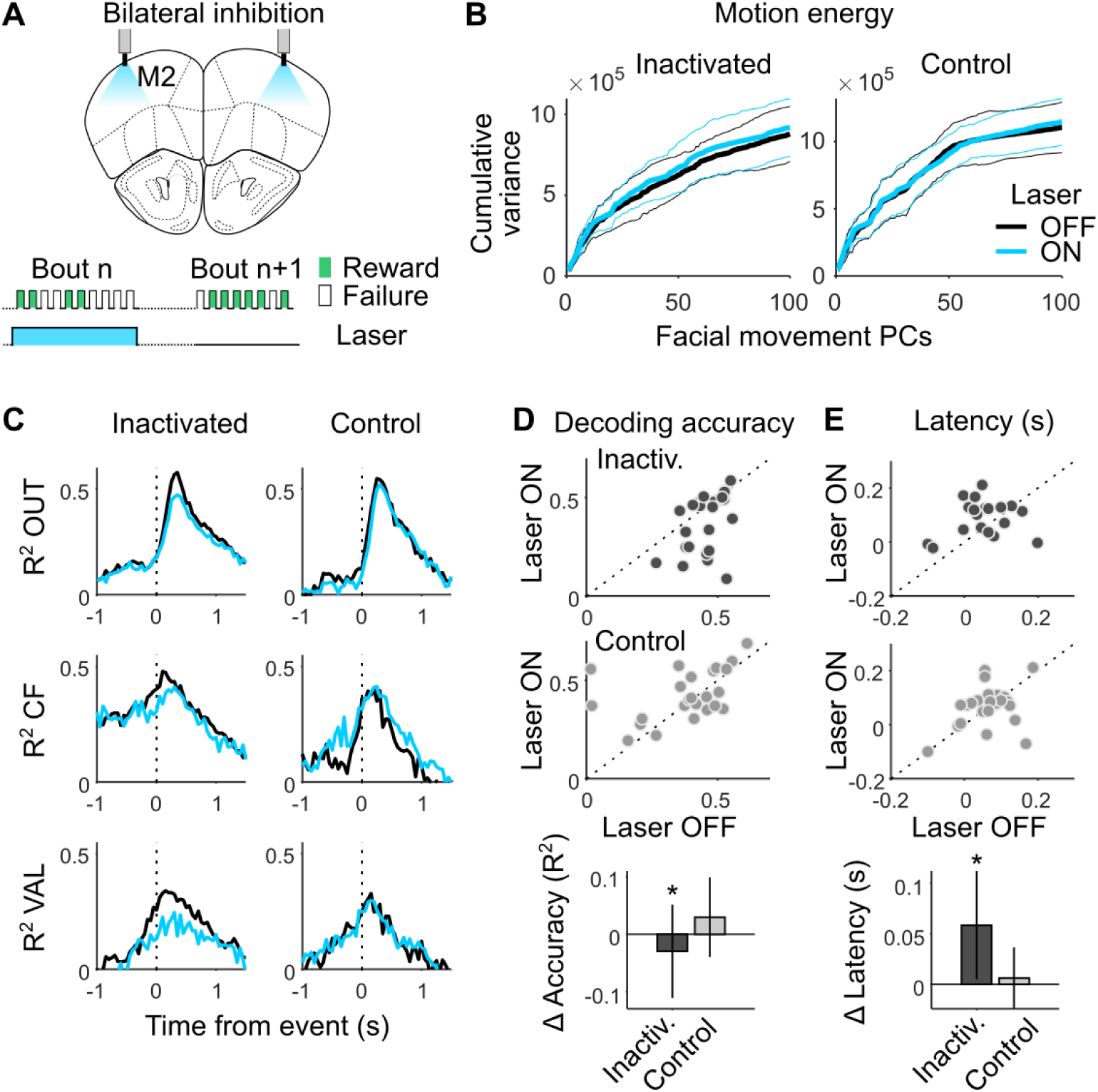
Effect of M2 inactivation on facial expressions of decision variables. A. Schematic representation of optic fiber placement on top of M2 (+2.5 anterior, ± 1.5 lateral of bregma). Bilateral photostimulation (5 mW power per fiber, 10 ms pulses at 75 Hz) was triggered by the first lick in 30% of trials and lasted until the last lick of the bout. B. Cumulative variance of the facial movement PCs during laser OFF (black) and laser ON (cyan) conditions in the inactivated group (5 mice, median ± m.a.d. across N = 24 sessions) and in the control group (3 mice, median ± m.a.d. across N = 24 sessions). C. Decoding accuracy of multivariate regression models predicting decision variables from facial movement PCs as a function of the time delay of the predictors during laser OFF (black) and laser ON (cyan) conditions (median across N = 24 sessions for both inactivated and control groups). D. Top & middle: Decoding accuracy in laser ON against laser OFF conditions. Dots below the identity correspond to representations of decision variables from facial movement PCs during laser ON being less accurate than that of the laser OFF condition. Bottom: Difference between the accuracy of laser ON and OFF conditions (median ± m.a.d. across 24 sessions for inactivated and control groups). One star indicates significance (p < 0.05) in the Wilcoxon signed-rank test. E. Same as in (D) but for decoding latency.

Once again, we used GLMs to decode the decision variables from facial movement PCs (Fig. S3) in laser ON condition (i.e., M2 silencing) and compared the latency and accuracy of the decodings to those observed in laser OFF condition (i.e., unperturbed M2 activity). While the variance of the facial movement PCs was unaffected by the photoinhibition (Fig. 5B, Wilcoxon signed rank test: p = 0.15 for inactivated mice; p = 0.28 for control mice), partial silencing of M2 decreased the accuracy by which decision variables could be predicted with facial movement PCs (Fig. 5C,D, ΔAccuracy inactivated mice: -0.03 ± 0.08, p = 0.018; ΔAccuracy control mice: 0.03 ± 0.07, p = 0.24; median ± m.a.d. across 24 sessions each, Wilcoxon signed rank test). Partial silencing of M2 also delayed the representation of decision variables from facial movement PCs (Fig. 5C,E, ΔLatency inactivated mice: 0.06 ± 0.05, p = 0.033; ΔLatency control mice: 0.006 ± 0.03, p = 0.43; median ± m.a.d. across 24 sessions each, Wilcoxon signed rank test). Altogether, these results suggest a direct role for M2 in generating facial expressions of decision variables.

## DISCUSSION

In this study, we investigated how facial movements in mice reflect ongoing brain computations during a foraging task. We found that there is a remarkable richness in the representational capacity of the face, which reflected not only the identity of the current behavioral strategy used by each mouse and the time-varying decision variable that underlies this strategy, but also, alternative time-varying decision variables that support strategies used during other epochs. The musculature of the face is thus a high-capacity information channel that tracks in real time a multiplicity of latent dynamical variables computed by the brain that are related in complex ways (e.g., temporal integration or reset) to immediately available external events, such as individual licks or their rewarding consequences. At a mechanistic level, we found that the facial expressions associated with these decision variables appear to originate from activity in the premotor cortex (M2) and may, in turn, influence activity in other cortical regions.

Traditional studies of the neural control of movement have focused on elucidating purposeful and adaptive behaviors, such as those required to perform a task (*25*). More recently, studies of seemingly meaningless expressions and movements of the face and body are also being linked to specific neural activity patterns (*15*, *24*). Establishing these relationships is important in at least two ways. The first, which has received more emphasis, relates to the potential ability of facial movements to explain neural activity that would otherwise be considered as “noise” or misattributed. The second, which we have focused on in this report, relates to their potential to be used as visible readouts of latent cognitive processes.

The face is a particularly interesting site of neural control in mammals, being well known as a locus of emotional expression. Emotional expressions, considered the canonical markers of internal states, are still quite mysterious despite their importance to human life. While traditional accounts identify a handful of emotional states, classification using statistical methods suggests a much richer variety (*7*, *26*). The discovery of facial expressions of emotions in mice (*9*, *27*) has opened the door for a better understanding of their neural basis. A key step in this direction has been to link facial expressions to controllable and quantifiable variables. Recent examples of this are facial expressions being linked to auditory-evoked responses (*13*, *14*) and to arousal state (*16*, *21*). Clayton et al. 2024 (*14*), in particular, showed that facial movement is not only more sensitive to the intensity of auditory stimuli, but also that it reports the spectral and temporal structure of sound. Their results however, point to a sophisticated but very fast transformation from sound to muscle-pattern, leaving open the extent to which the face can accurately report temporally extended cognitive variables associated with high-level cognition.

Decision-making tasks offer a powerful opportunity to link incidental expressions to temporally extended neural variables such as accumulated evidence. Previous studies showed that a running estimate of accumulated sensory evidence is present in the musculature of the arm before a choice is made in a decision making task where the arm is used to report the decision (*5*). Movement vigor has also been shown to report uncertainty in a foraging task in mice (nose-poking vigor; (*28*)) and decision confidence in a perceptual decision making task in humans (finger speed; (*29*)). The contents of working memory are also reflected in postural adjustment in rats (*3*, *4*). These results are consistent with accounts of computation which emphasize readiness in real-time (*30*), so that the state of the motor system continuously reflects the agent’s best guess of what to do at the moment.

In our study, we explored whether the cognitive significance of incidental facial expressions extends beyond (i) fast, reflex-like encoding of the current sensory environment (*13*, *14*) and (ii) the immediate demands that tasks requiring temporally-extended computations place on the motor system. To do so, we took advantage of the recent finding that mice compute several different decision variables simultaneously when performing a foraging task (*22*). This allowed us to search for correlates of internal variables that were unrelated (and indeed mathematically orthogonal) to the currently used behavioral strategy. Unexpectedly, we found that these non-task variables are expressed as much as task-related ones and that the face contains as much information about these decision variables as recordings from approximately a hundred neurons in the secondary motor cortex (M2).

The decision variables could be decoded from M2 activity earlier (approximately 50 ms) compared to facial movements. The delay between the representation of decision variables in M2 and in facial movement is consistent with previous work reporting that movement-related signals in the brain occurred at least 50 ms before detectable movement onset (*31*). Furthermore, partially inactivating M2 with optogenetics reduced our ability to decode decision variables from facial expressions (it increased the latency and reduced the accuracy of the decoding). Interestingly, Clayton et al., 2024 (*14*) found that inactivating the auditory cortex enhanced, rather than disrupted, the sensitivity of face to sound intensity, suggesting a diversity of cortical and subcortical pathways converging on the musculature of the face to report brain activity. Together, our findings suggest that M2 might be involved in generating the facial expressions associated with decision variables. In contrast, decoding decision variables from OFC and OC was only possible after their detection in facial expressions and with significantly lower accuracy compared to facial movements. This suggests that the presence of decision variables in the OFC and OC activity might be driven by proprioceptive feedback. Together, widespread motor signals related to efference copies, as well as driven by proprioceptive feedback, contribute to the prevalence of movement-related activity across various brain regions (*32*).

The ability of video and other digital readouts to disclose information that has heretofore been considered private or internal is rapidly being exposed. Unconscious patterns of gaze and head movements detected by virtual reality headsets can be used to rapidly reveal individual identity (*33*, *34*). Thus, understanding the limits of these technologies is important for shaping our views on the protection of privacy. Similarly to facial expressions associated with emotions (*9*), we found that the different decision-related expressions correlated with characteristic features of the face and appeared stereotyped across mice, potentially facilitating their recognition through generalizable methods. This, together with the remarkable expressivity of facial muscles – facial expressions surpassed ensembles of around one hundred M2 neurons in their ability to reveal both the decision variable guiding the animals’ choices and the alternative decision variables considered – suggests ample opportunities for non-invasive methods, like facial analysis, for inferring hidden brain computations. This highlights the need to devise effective regulatory policies to address the misuse of biometric technologies for privacy invasion (*35–37*).

## METHODS

### Behavioral approach

We briefly summarize experimental methods that have been described fully in (*22*). Mice (N = 16, male and female, 2-9 months old) were trained in a head-fixed version of a probabilistic foraging task. All experimental procedures were approved and performed in accordance with the Champalimaud Centre for the Unknown Ethics Committee guidelines and by the Portuguese Veterinary General Board (Direção Geral de Veterinária, approval 0421/000/000/2016).

Mice were first implanted with a head-bar under standard aseptic surgical procedures. After a recovery period (5 to 10 days), mice were water-restricted and only received sucrose water (10%) during the task. Mice were given 1 mL of water or 1 gram of hydrogel (Clear H2O) on days when no training or recording occurred or if they did not receive enough water during the task.

During the task, mice were head-fixed and placed on a linear treadmill. Running on the treadmill could activate the movement of two arms, which materialized two different foraging sites, via a coupling with digital Servo motors (Hitec HS-5625-MG). At the extremity of each arm, water flowed through a lick-port by gravity through water tubing and was controlled by calibrated solenoid valves (Lee Company). To lick at an arm and receive rewards, mice had to decrease their running speed for more than 250 ms below a threshold for movement (3 cm/s). Licks were detected in real time with a camera (FLIR Chameleon-USB3, 60 fps) located on the right side of the treadmill using a thresholding method in BONSAI (*38*). Each lick was rewarded in a probabilistic fashion by a small drop of water (1 μl) with a probability of 0.9 in the rewarding state. The small reward size ensured that there was no strong difference in licking rate between rewarded and unrewarded licks. Each lick could cause the state of the currently exploited site to transition from rewarding to unrewarding with a probability of 0.3. When such transition occurred, the currently unexploited site (the distal one) transitioned from an unrewarding state to a rewarding state. To leave the currently exploited site and reach the other one, mice had to restart running above the threshold for movement for more than 150 ms, and travel a fixed distance on the treadmill (around 16 cm).

### Neuropixels recording and processing

Here again, we briefly summarize electrophysiological methods that have been described fully in (*22*). Recordings (N = 10 recording sessions in 9 mice) were made using electrode arrays with 374 recording sites (Neuropixels “Phase3A”). Before each recording session, the shank of the probe was stained with red-fluorescent dye (DiI, ThermoFisher Vybrant V22885) to allow later track localization. Prior to the first recording session, mice were briefly anesthetized with isoflurane and administered a non-steroidal analgesic (Carprofen) before drilling one small craniotomy (1 mm diameter) over the secondary motor cortex (+2.5 mm anterior and +1.5 lateral relative to bregma) from either hemisphere. The craniotomy was cleaned with a sterile solution and covered with silicone sealant (Kwik-Sil, World Precision Instruments). After several hours of recovery, mice were placed in the behavioral setup and the probe was slowly advanced through the dura and slowly lowered to its final position (3 to 3.5 mm inside the brain). The probe was allowed to settle for at least 10 min before starting recording to ensure better stability of the recorded units. Recordings were acquired with SpikeGLX Neural recording system (https://billkarsh.github.io/SpikeGLX/) using the external reference setting and a gain of 500 for the AP band (300 Hz high-pass filter).

After the recording session, mice were deeply anesthetized with Ketamine/Xylazine and perfused with 4% paraformaldehyde. The brain was extracted and fixed for 24 hours in paraformaldehyde at 4 C, and then washed with 1% phosphate-buffered saline. The brain was sectioned at 50 μm, mounted on glass slides, and stained with 4’,6-diamidino-2-phenylindole (DAPI). Images were taken at 5x magnifications for each section using a Zeiss AxioImager at two different wavelengths (one for DAPI and one for DiI). To determine the trajectory of the probe and approximate the location of the recording sites, we used SHARP-Track (*39*). Characteristic physiological features were also used to refine the alignment procedure (especially the absence of spikes between frontal and olfactory cortical boundaries and the LFP signature in deep olfactory areas).

Neural data were first automatically spike-sorted with Kilosort2 (*40*) using MATLAB (MathWork, Natick, MA, USA). To remove artifacts, traces were “common-average referenced” by subtracting the median activity across all channels at each time point. Second, the data was manually curated using an open source neurophysiological data analysis package (Phy: https://github.com/kwikteam/phy). Units labeled as artifacts were discarded in further analyses. Units passing quality criteria (as described in (*22*)) were labeled as good and considered to reflect the spiking activity of a single neuron. For all analyses, otherwise noted, we averaged for each neuron the number of spikes into bins by considering a 200 ms window centered around each lick. Because the interval between each lick was on average around 150 ms, there was little overlap between two consecutive bins and each bin typically contained the number of spikes associated with only one single lick.

### Video monitoring and processing

Facial movements were monitored at 60 fps using an infrared camera (the FLIR Chameleon-USB3 for 56 sessions and the Sony PlayStation 3 Eye Camera for 2 sessions) positioned on the side of the animal. The videos were synchronized with the electrophysiological and the behavioral data for subsequent analysis. To extract movement signals from the videos, we used an open-source toolbox: FaceMap (*19*, *24*). Specifically, we performed SVD on the motion energy (the absolute value of the difference of two consecutive frames) to compute the 500 highest dimensions of video information. For each dimension, we averaged the signal in a 200 ms window centered around each lick, similar to the processing of the neural data.

To register the videos, 2D spatial masks corresponding to the facial PCs were aligned to a reference “mouse face”. Specifically, for each video, a set of control points (8) was positioned on the average face of the mouse for that session. These 8 points and the corresponding points on the reference face were used to estimate an affine transformation between the two images. The transformation was then applied to the facial masks in order to register each map to the same reference. The same transformation was also applied for each frame and session for the analysis where we concatenated videos from all sessions. To reconstruct the motion energy reflecting each decision variable, we weighed each facial mask by the corresponding model weights, and then performed a linear combination of all the weighted facial masks (N = 100).

### Photoinhibition

To optically silence M2 activity by stimulating ChR2 expressing VGAT-expressing GABAergic interneurons we used blue light from a 473 nm laser (LRS-0473-PFF-00800-03, Laserglow Technologies, Toronto, Canada, or DHOM-M-473-200, UltraLasers, Inc., Newmarket, Canada). Light was emitted from the laser through an optical fiber patch-cord (200 μm, 0.22 NA, Doric lenses), connected to a second fiber patch-cord with a rotatory joint (FRJ 1x1, Doric lenses), which in turn was connected to the chronically implanted optic fiber cannulas (M3 connector, Doric lenses). The power of the laser (5 mW) was calibrated before every session using an optical power meter kit (Digital Console with Slim Photodiode Sensor, PM100D, Thorlabs). The optical stimulation (10 ms pulses, 75 s^-1^, 5 mW) was turned on during 30% of randomly interleaved bouts. Light delivery started when the first lick was detected and was interrupted if the animal did not lick for 500 ms (which was in 98% of bouts after the last lick of the bouts).

### Data Analyses

All data analyses were performed with custom-written software using MATLAB. We used generalized linear regression models (GLMs) to fit the switch behavior (with GLM for Bernoulli responses) or the decision variables (all with GLMs for normal distributions) using facial movement PCs or neural activity as predictors. Model fits were performed on individual recording sessions (Fig. 2, 4 and 5) or on concatenated sessions when looking at strategies (Fig. 1 and 3). To avoid overfitting, we used nested cross-validation and elastic net regularization (*α* = 0.5) using the minimum lambda to select the hyper-parameter that minimizes prediction error. To assess the predictive power of the model, we also implemented a 5-fold cross-validation. Specifically, the model coefficients and hyperparameters were sequentially fit using a training set consisting of four-fifths of the data and the prediction was evaluated on the testing set consisting of the remaining one-fifth. The method was implemented until all the data had been used both for training and testing. The coefficients of determination (R^2^) reported as a metric of the goodness of fit were calculated from the cross-validated results.

To estimate the latency of decision variable representations in facial movement and neurons (Fig. 4 & 5), for each neuron and facial movement PC, we used a 200 ms sliding window with 75% overlap to create lagged versions of the binned activity. These lagged activity series were fed into separate, cross-validated, and regularized GLMs to predict the different decision variables at different delays. For each decision variable, we constructed a curve to depict how decoding accuracy changed with increasing delay across individual sessions. Latency of the decoding was then defined as the delay corresponding to the midpoint between the minimum and maximum decoding accuracy around the peak of the curve. We considered the median latency across the different decision variables to perform statistical analysis across sessions.

### LM-HMM

To detect the precise moment when animals switch between strategies within a session, we use a previously developed model that combines a framework based on hidden Markov models (HMM) with linear regression models (LM) (see (*22*)). The resulting ‘LM-HMM’ framework takes input-driven gaussian observations, modeling a time-varying linear dependence 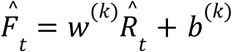 of normalized consecutive failures 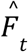 (observations) on normalized total rewards 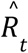 (inputs) across bouts *t=1,…T*; ɛ_*t*_ is i.i.d. gaussian noise with mean zero and variance σ^(*k*)^. For each session *m*, the normalized values 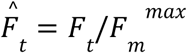 and 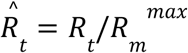 were obtained by min-maxing the raw values *F_t_*, *R_t_* on their within-session max 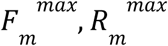, allowing us to fit a single model to all sessions where both inputs and observations were bounded between zero and one. Here, the slope *w*^(*k*)^, intercept *b*^(*k*)^ and noise variance σ^(*k*)^ depend on the hidden state *k=1,…,K*, each state representing a different strategy. Specifically, states with *w*^(*k*)^=0 or *w*^(k)^ >0 represent inference-based and stimulus-bound strategies, respectively. Bias *b*^(*k*)^ reflects persistent (large) or impulsive (small) behavior. We fit the model to all mice using an Expectation-Maximization algorithm to optimize parameters and we chose the best model complexity (3 states) using 3-fold cross-validation with maximum likelihood estimation and maximum a posteriori approaches (see (*22*) for details).

## ACKNOWLEDGEMENTS

We thank Joao P. Morais for support with behavioral training. This work was funded by CNRS (F.C.), Simons Foundation (F.C.: SCGB 969875; Z.F.M.: SCGB 543011), Marie-Curie postdoctoral fellowships (F.C.: HORIZON-MSCA-2021-PF-01 101062459; D.R.: HORIZON-MSCA-2021-PF-01 101063075), Fundação para a Ciência e Tecnologia (A.R.: LISBOA-01-0145-FEDER-032077 and PTDC/MED-NEU/4584/2021), the European Research Council Advanced Grant (Z.F.M.; 671251), and Champalimaud Foundation (A.R., Z.F.M.). This work was also supported by Portuguese national funds, through FCT - Fundação para a Ciência e a Tecnologia - in the context of the project UIDB/04443/2020 and by the research infrastructure CONGENTO, co-financed by Lisboa Regional Operational Programme (Lisboa2020), under the PORTUGAL 2020 Partnership Agreement, through the European Regional Development Fund (ERDF) and Fundação para a Ciência e Tecnologia (Portugal) under the projects LISBOA-01-0145-FEDER-02217 and LISBOA-01-0145-FEDER-022122.

## AUTHOR CONTRIBUTIONS

F.C., A.R. and Z.F.M. designed the study. F.C. and E.A. performed behavioral and optogenetic experiments. F.C. performed electrophysiological experiments and curated the data. D.R. processed the video data. F.C., D.R. and A.R. designed and performed the analyses. F.C., A.R and Z.F.M. wrote the manuscript. All authors reviewed the manuscript.

## SUPPLEMENTARY MATERIALS

**Supplementary Fig. 1.**
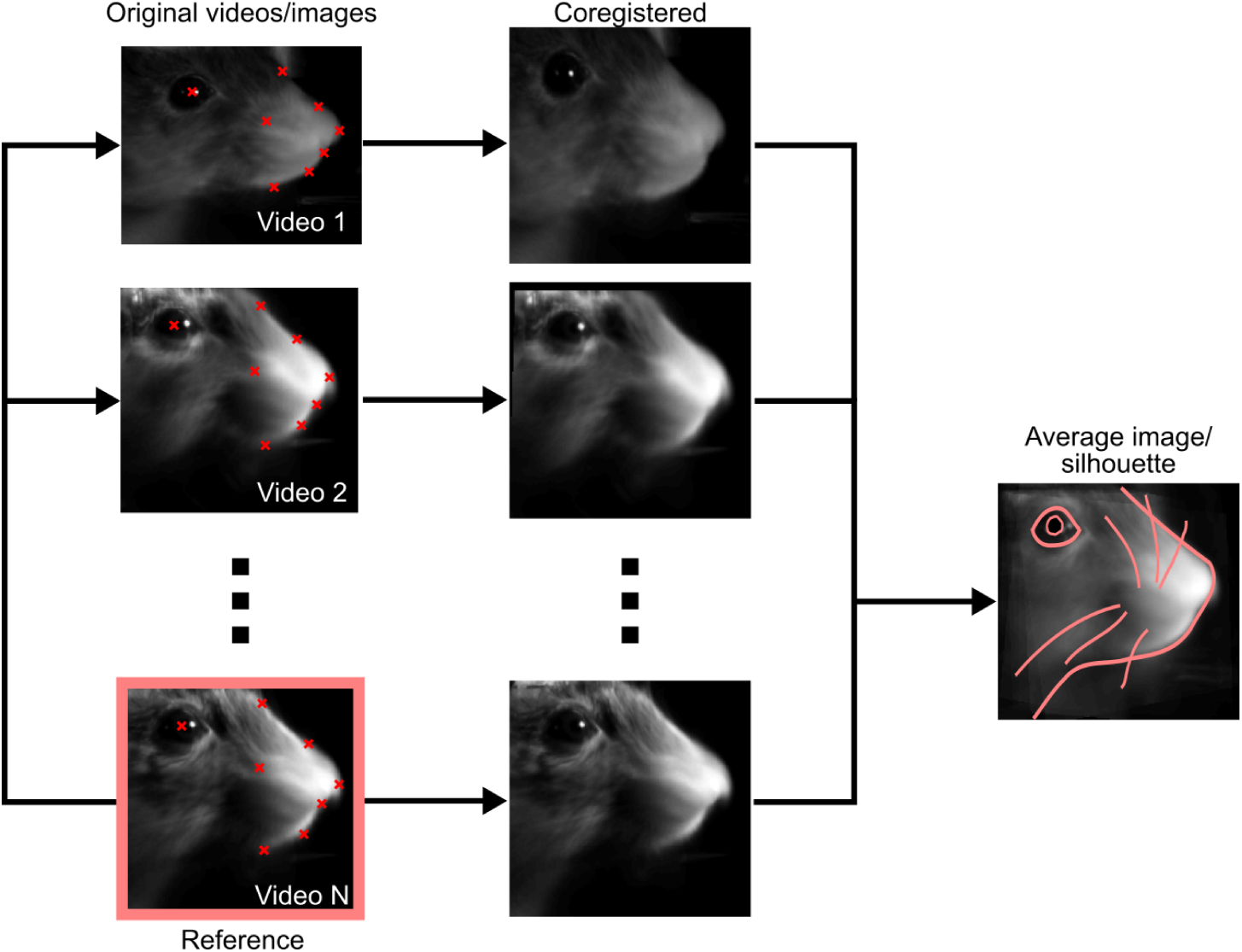
Coregistration of the face from different videos. An average frame for each video was used to label 8 distinct points on the mouse face (red labels). Those points were used to identify an affine transformation that could be used to co-register frames from each video to a reference one (pink, bottom) using *fitgeotrans* in MATLAB. The transformation was used to compare and average weighted maps (as in Fig. 2), concatenate videos (as in Fig. 3) and to define an average silhouette that was superimposed to images in Fig. 2.

**Supplementary Fig. 2.**
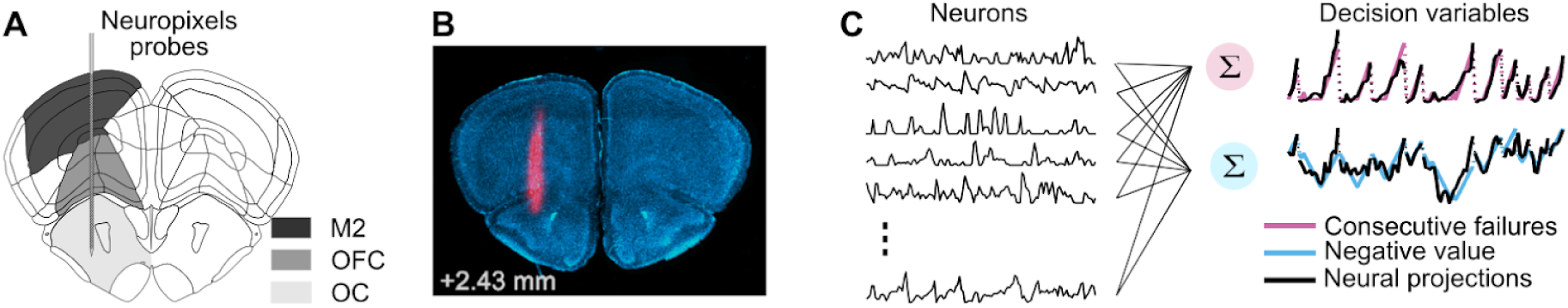
Decoding multiple cognitive variables from neural activity. **A.** We recorded with Neuropixels probes in multiple regions of the frontal cortex. Schematic target location of the neuropixels probe insertion. Vertical insertions were performed within a 1 mm diameter craniotomy centered around +2.5 mm anterior and +1.5 mm lateral from Bregma. **B.** An example of histology with the electrode track. We painted the probe with a pink, fluorescent dye to recover the probe’s location post-hoc. **C.** To decode the instantaneous value of multiple decision variables (pink & blue traces, right), we used regression models taking as predictors the activity of simultaneously recorded neurons in each brain region (black traces, left, example activity from M2). The model predictions (the weighted sums of neural activity, black trace right) overlap with the decision variables.

**Supplementary Fig. 3.**
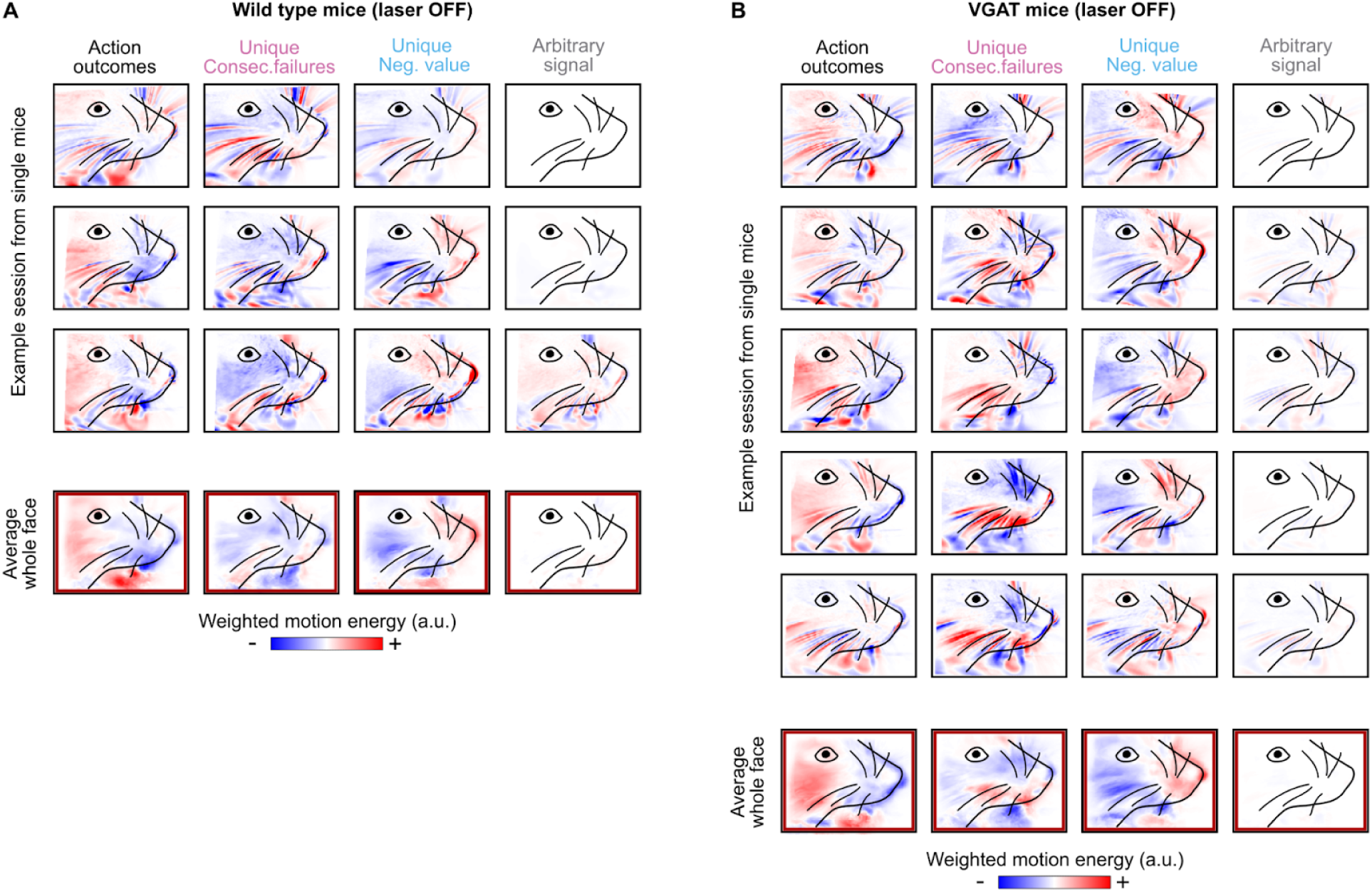
Stereotyped facial expressions of decision variables in wild-type and VGAT mice. **A.** Motion energy weighted by the models’ weights for a single example session (top) of each wild-type mouse (N = 3) and averaged across all sessions (N = 24, bottom) from all mice during the laser OFF condition. Red represents more movement than average, while blue indicates less movement than average. **B.** Same as in A but for a single example session (top) of each VGAT (N = 5) and averaged across all sessions (N = 24, bottom) from all mice during the laser OFF condition.

